# Modern Insights into Muscle Glycogen Phosphorylase Activity

**DOI:** 10.1101/2024.02.22.581477

**Authors:** Leonit Kiriaev, Jonathan S. Oakhill, Chrystal F. Tiong, Jane T. Seto, Vanessa G. Crossman, Kate G.R. Quinlan, Kathryn N. North, Peter J. Houweling, Naomi X.Y. Ling

## Abstract

Recent identification of new human muscle glycogen phosphorylation sites has renewed interest in understanding human variations in the regulation of glycogen metabolism and glucose homeostasis. This paper presents a detailed method for the measurement of glycogen phosphorylase (GPh) activity in skeletal muscle. Our approach incorporates modifications to existing radiolabelling assays, optimizing specificity and sensitivity while enabling the assessment of both active and total enzyme activity levels. The utilization of radioisotope tracers and scintillation counting ensures accurate quantification of GPh activity, which we use to validate a previously published reduction in GPh activity in an *Actn3* deficient mouse model. Moreover, we introduce a step-by-step guide for data acquisition, highlight the use of appropriate homogenization, discuss the need for allosteric activators/inhibitors and the importance of assay optimization to record a GPh activity assay for skeletal muscle. In conclusion, our refined method not only contributes to a deeper understanding of glycogen metabolism in muscle tissue but also provides a framework for future investigations, underscoring its role in advancing research on glycogen utilization and glucose homeostasis.

**NEW & NOTEWORTHY:** The study optimizes the glycogen phosphorylase radiolabelled activity assay, unveiling nuances in muscle homogenization, sample dilution, and caffeine inclusion. The research introduces standardized conditions, enhancing assay reliability and reproducibility across mouse strains to reveal sex specific variations in GPh activity and underscore novel distinctions in an Actn3 deficient mouse model. These findings advance our understanding of muscle glycogen metabolism, offering a crucial tool for researchers and facilitating meaningful inter-laboratory comparisons.

## INTRODUCTION

Glycogen, a branched polymer of glucose, serves as a crucial energy reserve in muscle tissue. Its metabolism is a fundamental process in energy homeostasis, playing a pivotal role in various physiological and pathological conditions, from exercise performance (1) to metabolic diseases such as obesity (2). Central to this intricate metabolic pathway is glycogen phosphorylase (GPh), the enzyme responsible for catalysing the breakdown of glycogen to release glucose-1-phosphate by phosphorolysis of the ⍰-1,4-glycosidic linkages in glycogen in the cytosol (Fig. 1A)(3).

**Figure 1:**
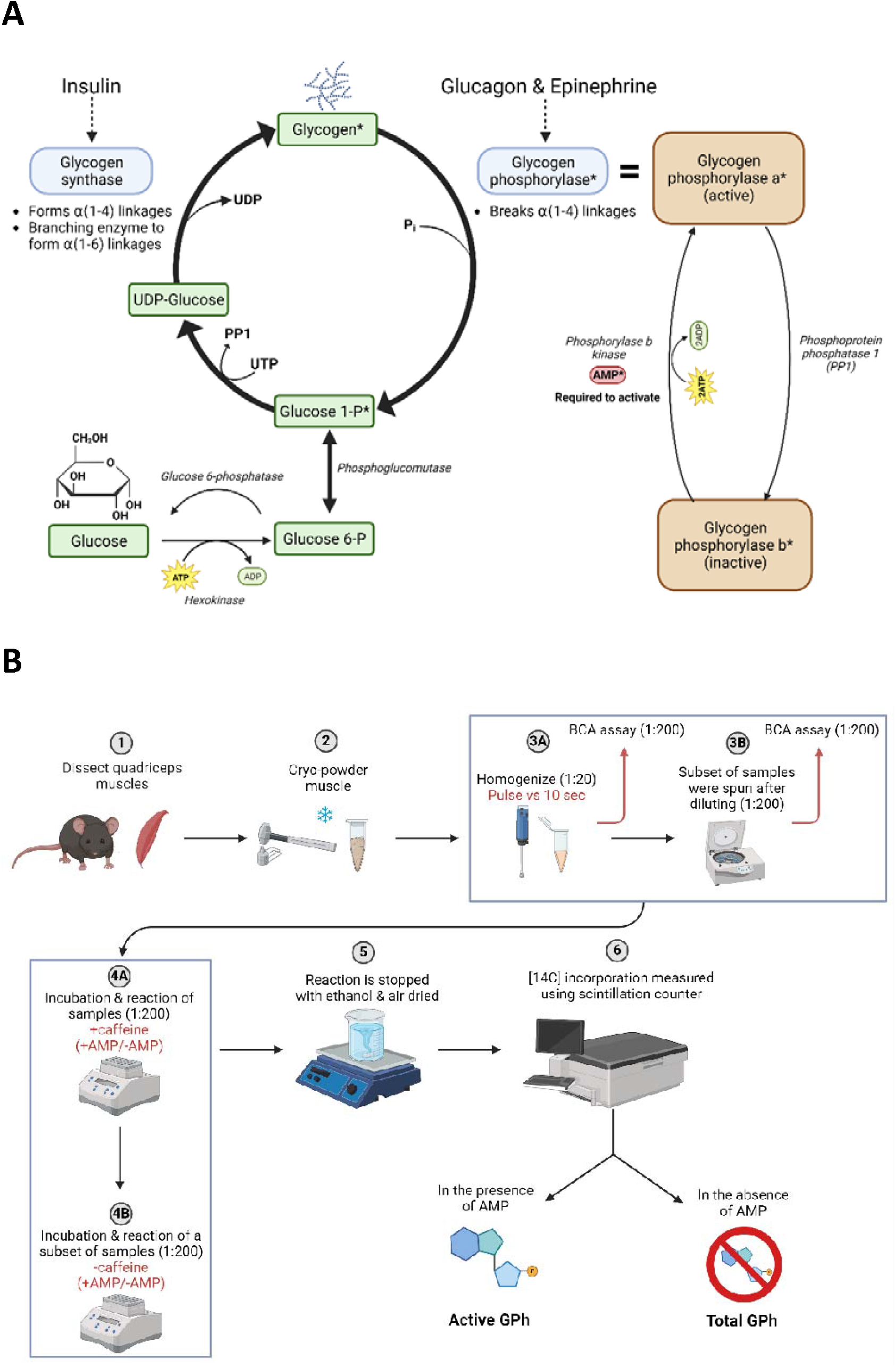
Biochemistry of glycogen metabolism and experiment flow. (A) Asterix (*) highlights key molecules involved in the glycogen phosphorylase activity assay. Corresponding workflow to measure GPh activity is shown in (B) highlighting variations in test conditions in red. Figures generated with BioRENDER.

Accurate assessment of GPh activity is a crucial aspect of understanding glycogen metabolism. To investigate these processes, reliable, precise and reproducible enzyme assays are essential. While the GPh activity assay is well-established using high performance liquid chromatography (4, 5), these enzyme assays with commercially available alternatives have low sensitivity, are complex and lack versatility in their application. While the assay is well-established for liver and brain tissue (5-7), its applicability to muscle, another significant site of glycogen storage, has been less explored.

Many past papers describing GPh activity radiolabelling assays in skeletal muscle lack detailed methodologies (8-12), often referring to an original methods paper from five decades ago (13). This trend suggests a reliance on established protocols without offering a standardized, nuanced understanding of the methodology’s application or potential refinements to optimize the assay for skeletal muscle.

Recent studies provide intriguing insights into the diverse activity levels reported by different laboratories. Manabe et al. (2013) reported GPh activity levels in skeletal muscle at approximately 70 nmol/mg (-AMP) and around 120 nmol/mg (+AMP), results that align with the findings of Testoni et al. (2017). In contrast, Quinlan et al. (2010) and the original GPh activity methods manuscript Gilboe et al. 1972 documented substantially higher active and total GPh activity values (more than threefold the level). This disparity in assay readouts underscores the necessity for standardized protocols and a detailed methodology that comprehensively determines assay sensitivity. Potential differences in homogenization techniques, extent of sample dilution for radiometric readout and the choice of whether to precipitate samples after homogenization are points for contention among methodologies. Additionally, the practice of correcting GPh activity radiometric counts per minute to protein levels (1, 10, 12) versus tissue weight (13-15) is inconsistently applied across literature. These discrepancies among laboratories underscore the need for standardized practices to ensure reliable and comparable results in the investigation of glycogen metabolism.

In the intricate landscape of glycogen phosphorylase modulation, allosteric inhibitors play a pivotal role in shaping enzymatic activity. Notably, caffeine, a well-established and natural GPh inhibitor, has been extensively studied (5, 16, 17). As the first natural compound proven to bind to the inhibitor site, caffeine has been a key focus in unravelling the regulatory mechanisms of GPh (18). In the context of GPh activity assays, caffeine is often employed to investigate the kinetics of GPh (19) and often used as a positive control in various inhibition studies (20, 21). Despite this, current GPh radiolabelling methodologies have considerable variability with a key divergence being the inclusion or exclusion of caffeine as an allosteric inhibitor (1). This variation introduces a critical challenge, as reported experimental conditions for assessing GPh activity widely differ among studies, posing difficulties in comparing results across laboratories. Beyond the selection of caffeine, discrepancies extend to enzyme sources encompassing isoforms from various species in both phosphorylated and dephosphorylated states (22). Furthermore, experimental conditions diverge in terms of enzyme concentration, substrate concentrations, assay temperatures, incubation times (ranging from no incubation to 15 minutes), buffer compositions and pH levels (17). This heterogeneity across methodologies underscores the inherent challenge in achieving standardized protocols, hindering the seamless comparison of GPh activity data and thus complicate our understanding of whether caffeine is required for the modulation of GPh enzyme activity.

This paper aims to address these critical gaps in the field by presenting a comprehensive study on optimizing the GPh radiolabelled activity assay. We delve into the intricacies of homogenization, sample dilution, and the inclusion of caffeine, aiming for a robust and sensitive assay that facilitates meaningful inter-laboratory comparisons and enhances our understanding of glycogen metabolism. Our previous work using this radiolabelling enzyme assay highlighted a 50% reduction in GPh activity in an α-actinin-3 deficient mouse model as an early and important driver towards a slower more oxidative muscle phenotype (1). While our previous investigations focused on comparing GPh activity between wild-type (WT) and *Actn3* knockout (KO) mice on the R129 strain, in this study, we specifically assess the consistency of these effects in *Actn3* mice on a C57BL/6 background. Through this holistic approach, our research not only contributes to refining GPh activity assessments but also provides a standardized methodology for studying this crucial enzyme in the context of skeletal muscle tissue.

## MATERIALS AND METHODS

### Animals and tissue collection

The *Actn3KO* mouse line, initially created in this laboratory by introducing a neo cassette knockout onto an R129 background (23), has also been backcrossed onto a C57BL/6 line and used to investigate the effects of *Actn3* deficiency in recent studies (24-27). For this study, homozygous wildtype (WT) and *Actn3* knockout (KO) mice were generated by het/het cross of *Actn3* mice on a C57BL/6 background. A total of 36 mice consisting of 10 KO males, 8 KO females and similar numbers of WT littermate controls were used for this study. All mice were fed food and water *ad libitum* and were maintained on a 12:12h cycle of light and dark.

Prior to muscle collection, 12-week-old mice were anaesthetised using 4% isoflurane delivered with oxygen and euthanized by cervical dislocation according to (MCRI AEC approved A760). Individual quadriceps muscles were then harvested and immediately snap frozen in liquid nitrogen and stored at -80°C for processing.

### Sample Preparation

Snap frozen quadricep muscles were cryogenically powdered using a metal socketed percussion mortar cooled via liquid nitrogen and placed on dry ice before being stored at -80°C. Approximately 10 mg of powdered muscle was used for homogenization, allowing the GPh activity assay to be performed on smaller muscle specimens for future preparations.

Samples were homogenized on ice for a continuous 10 second pulse using a polytron homogenizer in a buffer composed of 50 mM Tris-HCl (pH 7.4), 20 mM EDTA, 25 mM NaF, and 2 mM benzamidine. The homogenization was carried out using a homogenate-to-buffer ratio of 1:20 for Pierce BCA protein assay and 1:200 for GPh assay (unless otherwise specified). On a subset of 1:200 diluted samples, a spin step was performed after homogenization at 3000 rpm for 5 min at 4°C to test its effects on the assay readout (figure 1B).

### Glycogen Phosphorylase Assays

GPh catalyses the breakdown of glycogen to release glucose-1-phosphate, however, the enzyme is not specific to glycogen breakdown; it can also catalyse the reverse reaction, incorporating the addition of glucose-1-phosphate to the nonreducing end of a glycogen chain to extend the glycogen molecule. In this study, GPh assays were performed in the direction of glycogen synthesis to incorporate radiolabelled glucose-1-phosphate substrate into the growing glycogen chain. Notably, adaptations were introduced to the original methodology outlined by Gilboe et al 1972. The final assay components included at baseline 50 mM MES (pH 6.3), 7.5 mM [14C] glucose-1-phosphate (∼20 dpm/nmol), and 1.7 mg/mL glycogen (see table 1 below). The activity measurement of GPh enzyme in the absence of the allosteric activator AMP allows the level of the enzyme that is active (glycogen phosphorylase a, see figure 1) to be determined. This value can be compared with the activity measurement in the presence of 2.5 mM AMP, which stimulates the inactive portion of the enzyme (glycogen phosphorylase b) into activity and hence allows the total enzyme level to be assessed.

**Table 1:** Example of solution preparations for a final volume of 4.5 mL suitable for 25 samples.

| Reagent | Final concentration | Concentration in assay | Active GPh |  | Total GPh |
| --- | --- | --- | --- | --- | --- |
|  |  |  | (-AMP) Buffer (μL) | (-AMP) -caffeine Buffer (μL) | (+AMP) Buffer (μL) |
| 200 mM MES (Sigma, M3671) | 100 mM | 50 mM | 2250 | 2250 | 2250 |
| 300 mM Glucose-1-Phosphate (Sigma, G7000) | 15 mM | 7.5 mM | 225 | 225 | 225 |
| 50 mg/mL glycogen (Sigma, G0885) | 3.4 mg/mL | 1.7 mg/mL | 306 | 306 | 306 |
| 100 mM Caffeine in MQ (Sigma, C0750) | 0.5 mM | 0.25 mM | 22.5 |  |  |
| 100 mM AMP in MQ (Sigma, O1930) | 5 mM | 2.5 mM |  |  | 225 |
| 14[C]-Glucose-1-Phosphate (7 uL/mL) | 15000 dpm/50 μL | 15000 dpm/100 μL | 31.5 | 31.5 | 31.5 |
| dH2O |  |  | 1665 | 1687.5 | 1462.5 |

To elucidate the effects of the commonly used allosteric inhibitor (caffeine) on the GPh activity assay readout, the study included 0.25mM of caffeine in a subset of active GPh assays (figure 1B).

Optimized conditions were used for the final run consisting of n=8 samples per genotype and gender.

### Incubation and Reaction

Using an Eppendorf thermomixer, independent assays were incubated for various durations (0 minutes, 5 minutes, or 10 minutes) in different assay buffers: GPh assay buffer without AMP (-AMP), and GPh assay buffer with AMP (+AMP). Each of these assay conditions allowed the rate of glycogen synthesis to be assessed.

For each assay, 50 μL of the appropriate assay premix was placed in a 1.5 mL Eppendorf tube and equilibrated to 30°C. The assays were initiated by adding 50 μL of the muscle homogenate at the appropriate dilution (1:200). The reactions were incubated at 30°C and shaken at 1000 rpm for the specified assay duration.

### Assay Termination

To terminate the assays, 50 μL of the assay mixture was rapidly spotted onto a small square of labelled filter paper (2 cm x 2 cm), which rapidly absorbed the mixture within 1-2 seconds. The filter paper was then submerged into a large beaker containing 70% ethanol with a stirrer, allowing glycogen to precipitate onto the filter paper, halting the enzymatic reaction. The filter paper squares were left in 70% ethanol with continuous stirring for at least 30 minutes and then removed carefully to minimize any overlap and pinned onto a foamboard for air drying.

### Radioactivity Counting

The air-dried filter paper squares, now containing glycogen, were placed in scintillation vials containing 4 mL of Ultima Gold XR scintillant and vortexed for 30 seconds to mix. The radioactivity associated with the 14C label incorporated into glycogen on each filter paper disc was quantified using a scintillation counter. Additionally, aliquots of each assay buffer were placed in separate scintillation vials and counted to determine the disintegrations per minute (dpm/nmol) of [14C] glucose-1-phosphate.

TriCarb^®^ 4810 TR Liquid Scintillation Analyzer (PerkinElmer) was calibrated and normalized according to the manufacturer’s instructions before assay samples were counted. For each count, the following no muscle controls were implemented in duplicate as background control: scintillation fluid only, - AMP buffer (10 μL), +AMP buffer (10 μL), -AMP buffer (50 μL), +AMP buffer (50 μL).

The counts obtained from each assay were used to calculate the rate of [14C] incorporation into glycogen by GPh in nmol/min.

### Protein Content Measurement

The total protein content of the muscle homogenates was quantified using the Pierce BCA Protein Assay (23225) following the manufacturer’s instructions. Bovine serum albumin (BSA) standards were used for calibration to generate a 6-point standard curve in triplicates, and homogenates were assayed in duplicates. The protein content, expressed in milligrams (mg) of protein per extract, was used to normalize the activities of each enzyme from nmol/min to specific activities (nmol/min/mg protein).

### Assay Optimization

Prior to conducting experiments with experimental samples, the assays were optimized to ensure linearity with respect to the amount of extract added, incubation time, extent of homogenization, and absence/inclusion of caffeine. Optimal volumes of extract and time-points were selected based on these optimizations.

### Statistical analyses

Data are presented as means +/- standard deviation. Paired t-tests were used when different conditions were tested upon the same sample. Unpaired t-tests were used when comparisons were made between two groups (WT and KO) with both conducted using a significance threshold of 0.05. Statistical differences displayed within graphs: * *P*<0.05; \*\**P*<0.01; \*\*\**P*<0.001. All statistical tests and curve fitting by linear regression were performed using the statistical software package Prism (version 9.1.1; GraphPad).

## RESULTS

GPh radiolabelling assays were conducted to investigate GPh enzymatic activity and its role in glycogen synthesis, where it catalyses the formation of glycogen from glucose molecules. We assayed GPh activity in WT and KO muscles in the absence of the allosteric activator AMP, which allows the level of enzyme that is active to be determined. This value can then be compared with the activity measurement in the presence of AMP, which stimulates the inactive portion of the enzyme into activity, and hence allow the total GPh enzyme level to be assessed.

Extensive homogenization for at least 10 seconds produced more uniform and consistent protein samples to be used for the assay as shown by figure 2A, where there is a reduction in the differences between points for both active GPh activity (WT 119.7 to 2.9/KO 220.9 to 48.2 nmol/min/mg) and total GPh activity (WT 272.7 to 142.3/KO 489.6 to 62.7 nmol/min/mg). The same assay was performed in the presence of 0.25 mM of caffeine in Figure 2B but this had no further effect on GPh activity.

**Figure 2:**
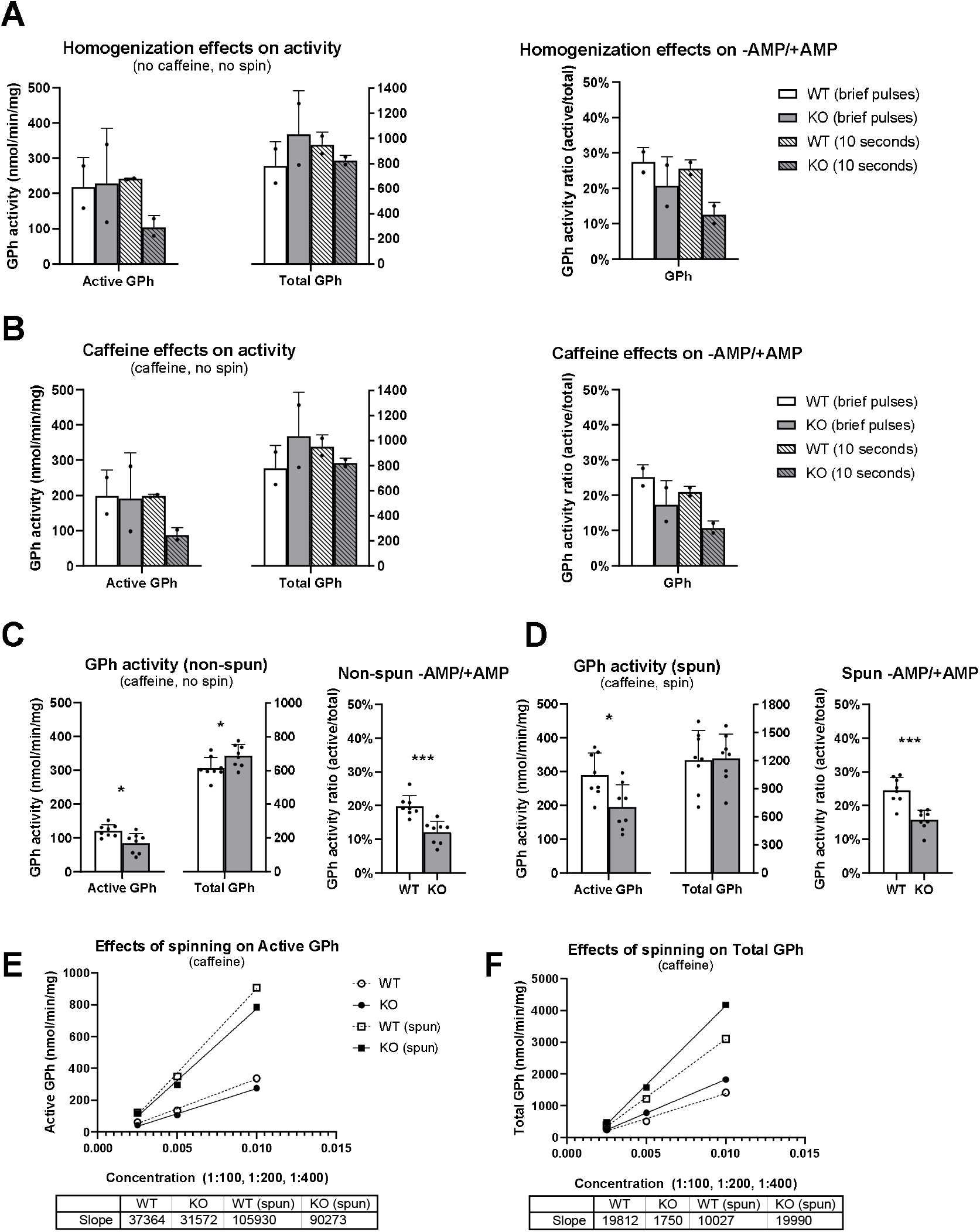
GPh assay optimization. Assay yield was optimized with respect to (A) homogenization time in the absence of caffeine for either 5 brief pulses or 10 seconds of continuous blending and (B) inclusion of caffeine on both homogenization conditions (n=2 male quads per genotype). For all bar graphs, active and total GPh levels are on the left and GPh activity ratio (Active/Total) on the right. With optimized conditions (10 seconds homogenization and caffeine), the assay was used to assess a larger cohort of tissues (n=8 male quads per genotype) in the (C) absence and (D) inclusion of spin after homogenization with a standard curve generated for a single titrated sample for each genotype at 1:100, 1:200 and 1:400 dilution. The (E) active GPh readout and (F) total GPh readouts for these titrated samples corrected for protein concentration are shown. See supplementary figure 1 for readouts for 2E and 2F not corrected for protein concentration.

Following homogenization, samples that were spun at 3000 rpm for 5 minutes (Figure 2D) showed an increase in both active and total GPh readout, as well as an increase in WT and KO -AMP/+AMP GPh activity ratio, compared to samples not spun (Figure 2C). This corresponded to increases in both active (Figure 2E) and total (Figure 2F) GPh activity across a single test sample with assays performed at 1:100, 1:200 and 1:400 dilutions. BCA protein concentration readouts for 2C and D are shown in supplementary figure 1A for reference.

Using optimized GPh activity assay conditions, the radioactive counts per minute (CPM) for 0, 5 and 10 minute incubation times show that assay sensitivity was not lost at higher (1:100) and lower (1:400) protein concentrations in the absence (Figure 3A) and presence of AMP (Figure 3B). When the gradient across these points were plotted out in subsequent graphs to assess the linearity for WT and KO samples, an R^2^ close to 1 is seen for both male (Figures 3C and D) and female (Figures 3E and F) mouse muscle samples for both active and total GPh activity.

**Figure 3:**
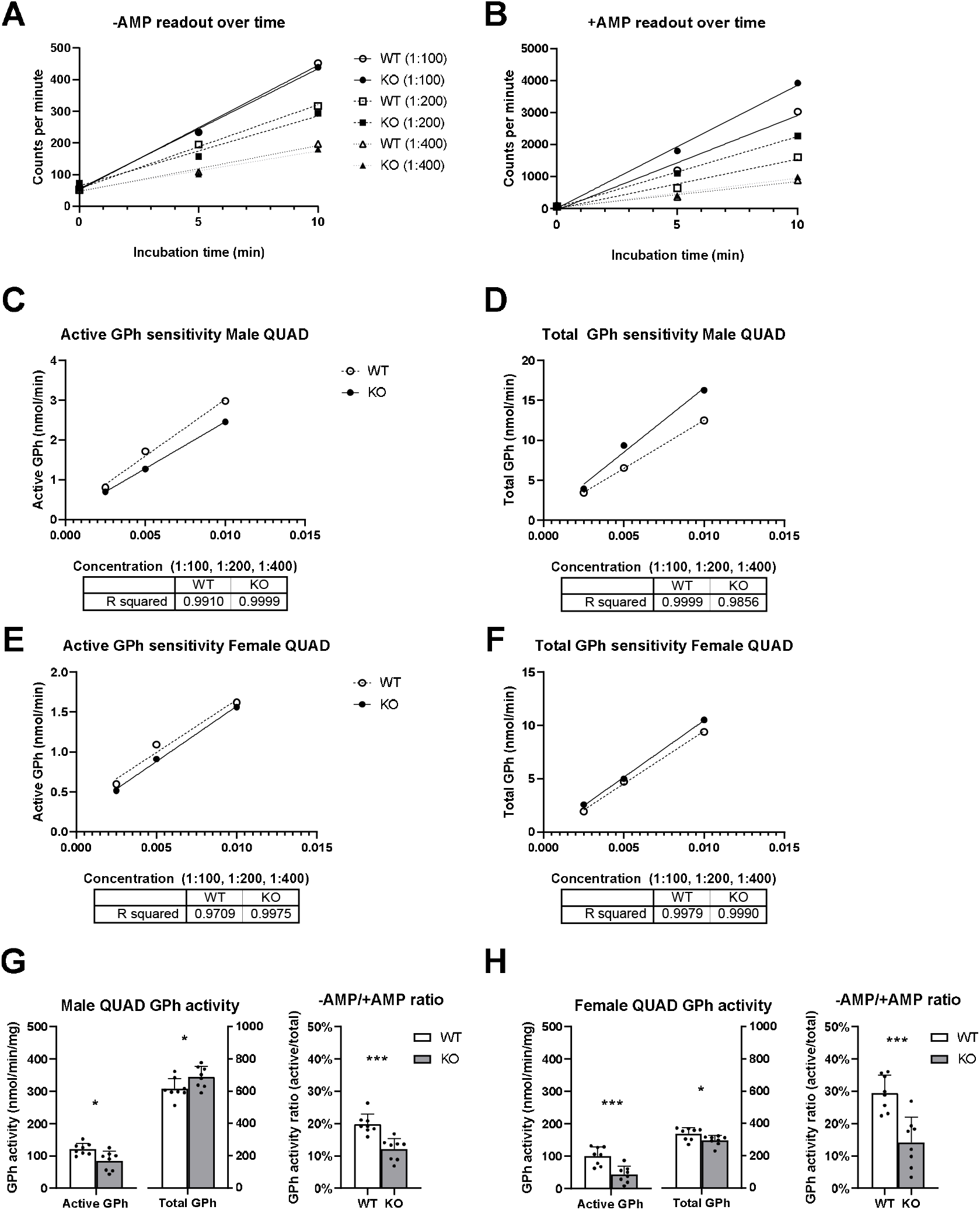
Assay sensitivity across different protein concentrations. Using optimized conditions (10 seconds homogenization, samples not spun and caffeine included) sensitivity of the assay was assessed at 1:100, 1:200 and 1:400 protein dilution on a single WT and KO quadriceps sample from male and female mice. A) Shows the radioactive count per minute readout for 0, 5 and 10min incubation times in the absence of AMP (active) and presence B) of AMP (total) in male WT and KO quadriceps muscles. For each of these experiments a 3-point standard curve (not corrected for protein loaded) was generated at 1:100, 1:200, 1:400 dilution using the slope of these graphs to assess the linearity of the assay for (C) active and (D) total GPh. The same was replicated in female quadriceps in (E) for active and (F) for total. Using optimized conditions and a protein dilution of 1:200, the active GPh activity, total GPh activity and GPh activity ratio (active/total) are shown for (G) male quadricep muscles (n=8 per genotype) and (H) female quadricep muscles (n=9 per genotype).

Finally, using a full cohort of muscle samples harvested at the same timepoint, we observed notable sex differences in GPh activity within both WT and KO mice cohorts. Both male and female WT mice show consistent active GPh activity readouts, however, female mice exhibited lower total GPh activity compared to males (335.4 compared to 615.1 nmol/min/mg) which resulted in a higher activity ratio (29.5%) compared to male counterparts (19.8%).

In KO mice, males demonstrated a 30% reduction in active GPh activity compared to WT and a 11.7% increase in total GPh activity which corresponded to a 7% reduction in activity ratio (-AMP/+AMP) (Figure 3G). When repeated on female KO mouse samples in Figure 3H, we demonstrated a 57% reduction in active GPh activity compared to WT and 12% decrease in total GPh activity which corresponded to a 15% reduction in activity ratio (2 fold change in -AMP/+AMP).

## DISCUSSION

Numerous considerations come into play when optimizing an enzyme activity assay. These encompass selecting and fine-tuning the composition of the buffer, determining the appropriate type and concentration of the enzyme, specifying the substrate type and concentrations, establishing optimal reaction conditions, and choosing the suitable assay technology. Here, we discuss a series of important methodology steps that we have identified within our study that will impact the optimal reaction conditions to perform the radiolabelled GPh activity assay.

### Importance of extensive muscle homogenization

In this study, muscle homogenization played a crucial role in the success and accuracy of the GPh radiolabelling assay. Traditional techniques involving short ‘bursts’ or brief ‘pulses’ are designed to isolate myofibrillar proteins while minimising denaturation (28). However, as depicted in Figure 2A, the process of homogenization using a longer continuous pulse on ice for 10 seconds compared to brief pulses is pivotal in preparing muscle tissue samples for the assay in order to ensure sample consistency across all the parameters tested, while ensuring cellular components such as GPh were released with minimal enzyme degradation at a 1:20 dilution. In pilot experiments, the same procedure at higher protein concentrations or shorter bursts of homogenization at the same concentration were attempted, both scenarios left pieces of the muscle bolus suspended in solution with low GPh activity across all samples. This emphasises the importance of tailoring the homogenization in order to ensure maximum release of cellular constituents and GPh activity yield.

### Inclusion of caffeine did not impact assay yield in the absence of AMP

Caffeine, a well-known methylxanthine compound, is increasingly gaining attention in the context of glycogen metabolism and its possible role as a regulator of GPh activity. Recent studies suggest that caffeine could influence the phosphorylation status of GPh and, in turn, affect glycogen breakdown (5, 16, 17). When we integrated caffeine into the GPh activity assay without altering any other parameters for active GPh, as depicted in Figure 2B, it offered us a unique opportunity to delve into the ability of caffeine to bind to the GPh inhibitor site to alter the conformation of the enzyme (18). In our assays, the absence (Figure 2A) or presence (Figure 2B) of caffeine did not significantly alter GPh activity in WT and KO QUAD muscles. Our results may be attributed to the absence of AMP in both test scenarios. AMP serves as an allosteric activator of glycogen phosphorylase b, changing its conformation and activating it (29). In its absence, the inactivation of AMP-activated protein kinase phosphorylase b kinase dictates the enzyme assay readout more than the presence/absence of caffeine (Figure 1A) rendering its effects negligible in other studies utilizing the same protocol (1). Another possibility is the employment of supraphysiological caffeine concentrations as discussed by Rush et al. 2014, where previous studies (30) have likewise shown caffeine did not displace the AMP activated form of phosphorylase a, thereby having a minimal effect in our experiment.

### Spinning post homogenization influences GPh readouts corrected for protein concentration

Conventionally, centrifugation is utilized to separate the particulate fraction (e.g., cellular debris) from the soluble fraction of the homogenate. Often this step is carried out to ensure a clear and homogenous solution that facilitates enzyme assays, particularly for soluble GPh which should be present in the supernatant post spinning. In our study, spinning down samples had no effect on the GPh readout of uncorrected radiolabelled counts per minute (Supplementary figure 1B and 1C), but affects total protein concentrations (Supplementary figure 1A), with significantly lower protein concentrations in the supernatant of the spun samples (WT 84-141μg/mL and KO 93-112μg/mL) compared to the non-spun equivalents (WT 218-335μg/mL and KO 89-252μg/mL). This resulted in significantly higher GPh activity in spun compared to non-spun samples when corrected for total protein, as shown in Figures 2C and D. As illustrated in Figures 2E and F, spinning samples increases the readout by 2-3 fold for both active and total GPh activity.

Although spinning down samples increases the magnitude of GPh readings following protein concentration correction, not spinning samples may provide the more accurate GPh readout. Firstly, not spinning the samples maintains the physiological relevance of the samples. Muscle tissue, particularly in its native state, contains a complex microenvironment where enzymes and substrates are intricately organized. Spinning down homogenates could disrupt this natural organization, potentially leading to an overestimation of GPh activity in the soluble fraction, as shown in Figure 2C and D. By omitting the centrifugation step, our assay reflects the native conditions more accurately.

Secondly, not centrifuging muscle homogenates minimizes the loss of important cellular components and substrates that may be present in the particulate fraction. This could be particularly relevant in the context of the GPh assay, where the enzyme’s interaction with its heavier polymer substrate (glycogen) and modulators is crucial for accurate measurements.

### Detection of sex-specific differences in GPh activity

The application of the GPh activity assay to a full cohort of muscle samples harvested simultaneously provides insights into notable gender-specific differences in both the WT and α-actinin-3 deficient mice cohorts (31). Here we show that in WT mice, female quadriceps muscles have higher GPh enzyme levels that correspond to a higher GPh activity ratio (-AMP/+AMP) compared to male counterparts. These discernible differences underscore the significance of considering sex-related factors in studies pertaining to glycogen metabolism.

Importantly, one of the primary phenotypes associated with α-actinin-3 deficiency is an alteration in skeletal muscle metabolism, with a shift towards a slower more oxidative phenotype. Our previous work highlights a ∼50% reduction in GPh activity (1) as an early and important driver of this phenotype in *Actn3* KO mice on an R129 background. Here, we have again demonstrated a reduction in GPh activity in KO mice on a C57BL/6 background.

Compared to WT, male *Actn3 KO* mice (Figure 3G) showed a significant 30% reduction in active GPh activity and an 11.7% increase in total GPh activity, resulting in a 7% reduction in the activity ratio (- AMP/+AMP). This discrepancy in active GPh activity and the corresponding alteration in the activity ratio suggests a nuanced response to AMP modulation in male mice. The genotypic differences of GPh activity in female mouse samples (Figure 3H) are more pronounced, with KO showing 57% reduction in active GPh activity, a 12% decrease in total GPh activity, and a resultant 15% reduction in the activity ratio (-AMP/+AMP) compared to WT. These gender-specific variations in GPh activity highlight the importance of considering sex-related factors in studies involving glycogen metabolism and emphasize the utility of our assay in elucidating such nuanced differences. The assay’s sensitivity across different protein concentrations and its ability to discern gender-specific variations underscore its applicability and reliability in investigating complex metabolic processes, offering valuable insights into the regulation of GPh activity in diverse physiological contexts.

### Sensitivity and optimized assay conditions

The results of our study demonstrate the robustness and sensitivity of the GPh activity assay under varying protein concentrations, providing confidence in its reliability for diverse experimental conditions. Notably, the radioactive counts per minute for 0, 5, and 10-minute incubation times revealed that the assay maintains its sensitivity at both higher (1:100) and lower (1:400) protein concentrations, both in the absence (Figure 3A) and presence of AMP (Figure 3B). The consistent linearity observed across a range of protein concentrations, as depicted in previous graphs for both male (Figures 3C and D) and female (Figures 3E and F) mouse muscle samples, underscores the ability of the assay to accurately quantify active and total GPh activity.

Our study has provided valuable insights into optimizing the sensitivity of the GPh activity assay, thereby enhancing its utility for investigating muscle glycogen metabolism. Our findings suggest that achieving the ideal sensitivity for the assay primarily relies on the thorough homogenization of muscle samples, approximately 10 seconds, to ensure the complete breakdown of muscle tissue and the release of GPh.

Moreover, our research indicates that sample dilution is a critical consideration for assay sensitivity. We found that dilution ratios of 1:100 or 1:200 can provide robust readouts without compromising sensitivity, ensuring that GPh activity measurements are accurate across a range of sample concentrations. These findings underscore the importance of finding the right balance between dilution and sensitivity to obtain meaningful and reproducible results.

The inclusion of a spin step and caffeine in the assay remains a subject of debate. Our data suggests that omitting the spinning step is advantageous for preserving the physiological relevance of the samples, and the role of caffeine in modulating GPh activity warrants further investigation. These findings provide researchers with a nuanced perspective on assay optimization, offering flexibility in tailoring the methodology to specific research objectives.

In addition to the optimization strategies discussed, it is important to highlight the robustness of the GPh activity assay in our study. Our findings have demonstrated that the assay is not only sensitive but also highly reproducible and reliable when applied to muscle samples across different mouse strains performed by different researchers. The ability to consistently measure GPh activity in muscle samples further enhances the assay’s versatility and relevance in physiological and pathological contexts where muscle glycogen metabolism plays a crucial role. By confirming the reproducibility and reliability of the assay for muscle, we provide researchers with a valuable tool for gaining deeper insights into the complex dynamics of muscle glycogen metabolism and the regulation of GPh in these tissues.

The lack of commercially available GPh kits in the market that are sensitive enough to be applied to skeletal muscle have prompted the need for customized and optimized protocols, as detailed in our research. Notably, the assay’s relevance extends beyond basic convenience; it directly responds to potential findings such as the identification of new GPh phosphorylation sites in humans and a recent analysis of the UK Biobank linking the presence of *ACTN3* with an increase in obesity related phenotypes (32).

In conclusion, our study’s comprehensive evaluation of the GPh activity assay highlights its adaptability and dependability when applied to muscle tissues, expanding its traditional use beyond liver and brain samples. The assay’s sensitivity, combined with its reproducibility and reliability, positions it as a valuable resource for a broader range of research questions in the fields of physiology, metabolism, and disease.

## AUTHOR CONTRIBUTIONS

L.K., J.O., C.T., J.S., K.Q., K.N.N., P.J.H., N.X.Y.L

L.K., C.T., J.S., N.X.Y.L generated data for the study.

L.K., P.J.H., N.X.Y.L analysed and interpreted the results.

L.K. wrote the manuscript and all authors reviewed the manuscript.

K.Q., K.N.N., P.J.H., N.X.Y.L conceptualized the study.

## SUPPLEMENTARY FIGURES

**Figure 1:**
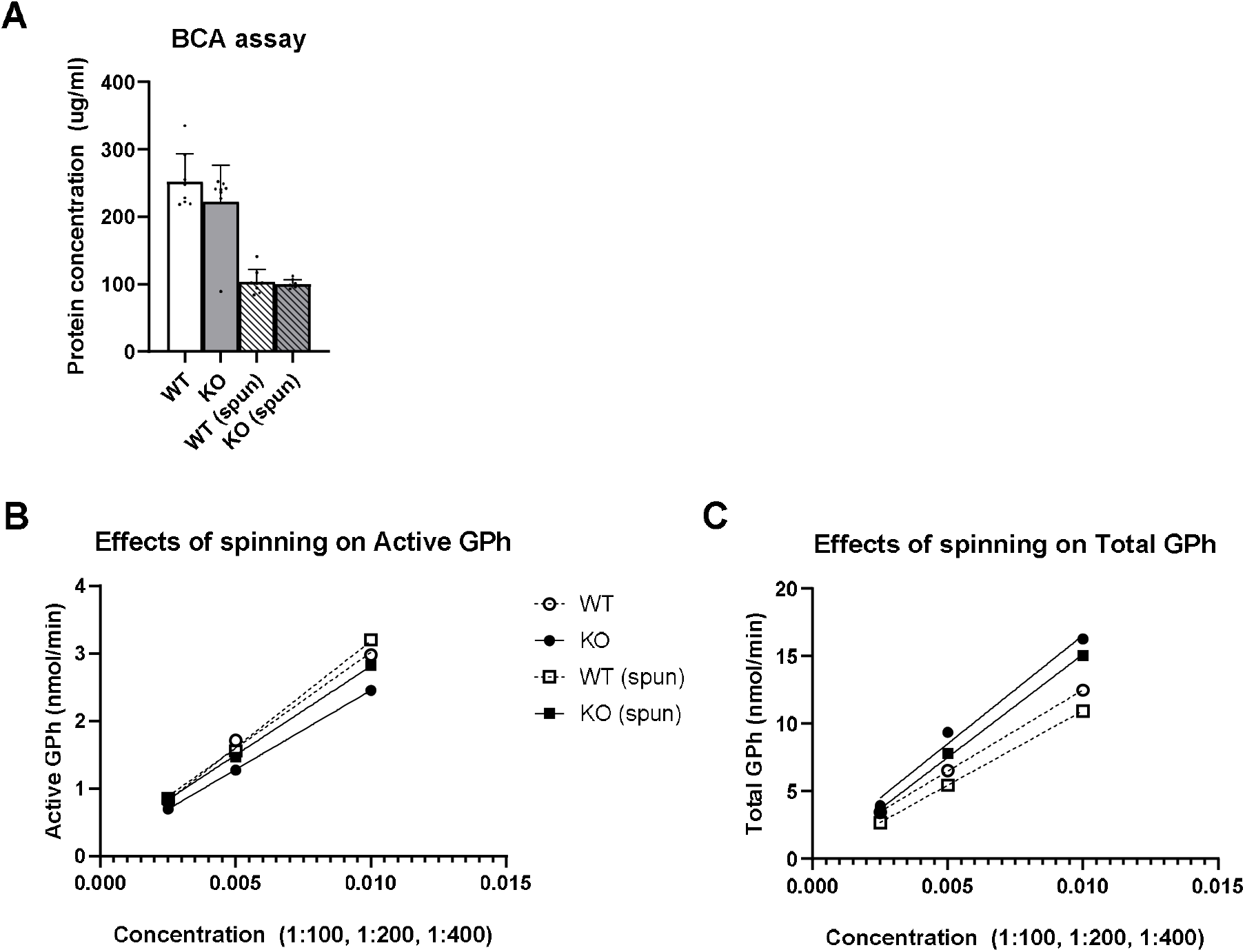
Protein concentration (A) determined by BCA protein assay for male quads samples in Figures 2C and D prepared without and with a spin step. BCA protein assays were conducted with 25μL of 1:200 homogenized extract. Raw readouts of radioactive labelled [14C] incorporated into glycogen by GPh for 2E and 2F not corrected for protein concentration are shown in (B) and (C) respectively.

## REFERENCES

1. Quinlan KGR, Seto JT, Turner N, Vandebrouck A, Floetenmeyer M, Macarthur DG, Raftery JM, Lek M, Yang N, Parton RG, Cooney GJ, and North KN. α-Actinin-3 deficiency results in reduced glycogen phosphorylase activity and altered calcium handling in skeletal muscle. Hum Mol Genet 19: 1335–1346, 2010.

2. Lu B, Bridges D, Yang Y, Fisher K, Cheng A, Chang L, Meng Z-X, Lin JD, Downes M, Yu RT, Liddle C, Evans RM, and Saltiel AR. Metabolic Crosstalk: Molecular Links Between Glycogen and Lipid Metabolism in Obesity. Diabetes 63: 2935–2948, 2014.

3. Adeva-Andany MM, González-Lucán M, Donapetry-García C, Fernández-Fernández C, and Ameneiros-Rodríguez E. Glycogen metabolism in humans. BBA Clin 5: 85–100, 2016.

4. Nakamura M, Makino Y, Takagi C, Yamagaki T, and Sato M. Probing the catalytic site of rabbit muscle glycogen phosphorylase using a series of specifically modified maltohexaose derivatives. Glycoconj J 34: 563–574, 2017.

5. Szabó K, Kandra L, and Gyémánt G. Studies on the reversible enzyme reaction of rabbit muscle glycogen phosphorylase b using isothermal titration calorimetry. Carbohydr Res 477: 58–65, 2019.

6. Mathieu C, Li de la Sierra-Gallay I, Duval R, Xu X, Cocaign A, Léger T, Woffendin G, Camadro JM, Etchebest C, Haouz A, Dupret JM, and Rodrigues-Lima F. Insights into Brain Glycogen Metabolism: THE STRUCTURE OF HUMAN BRAIN GLYCOGEN PHOSPHORYLASE. J Biol Chem 291: 18072–18083, 2016.

7. Zhu Y, Fan Z, Zhao Q, Li J, Cai G, Wang R, Liang Y, Lu N, Kang J, Luo D, Tao H, Li Y, Huang J, and Wu S. Brain-Type Glycogen Phosphorylase Is Crucial for Astrocytic Glycogen Accumulation in Chronic Social Defeat Stress-Induced Depression in Mice. Front Mol Neurosci 14: 819440, 2021.

8. Irimia JM, Rovira J, Nielsen JN, Guerrero M, Wojtaszewski JF, and Cussó R. Hexokinase 2, glycogen synthase and phosphorylase play a key role in muscle glycogen supercompensation. PLoS One 7: e42453, 2012.

9. Jindal HK, Merchant E, Balschi JA, Zhangand Y, and Koren G. Proteomic analyses of transgenic LQT1 and LQT2 rabbit hearts elucidate an increase in expression and activity of energy producing enzymes. J Proteomics 75: 5254–5265, 2012.

10. Manabe Y, Gollisch KS, Holton L, Kim YB, Brandauer J, Fujii NL, Hirshman MF, and Goodyear LJ. Exercise training-induced adaptations associated with increases in skeletal muscle glycogen content. FEBS J 280: 916–926, 2013.

11. Montori-Grau M, Minor R, Lerin C, Allard J, Garcia-Martinez C, de Cabo R, and Gómez-Foix AM. Effects of aging and calorie restriction on rat skeletal muscle glycogen synthase and glycogen phosphorylase. Exp Gerontol 44: 426–433, 2009.

12. Testoni G, Duran J, García-Rocha M, Vilaplana F, Serrano AL, Sebastián D, López-Soldado I, Sullivan MA, Slebe F, Vilaseca M, Muñoz-Cánoves P, and Guinovart JJ. Lack of Glycogenin Causes Glycogen Accumulation and Muscle Function Impairment. Cell Metab 26: 256–266.e254, 2017.

13. Gilboe DP, Larson KL, and Nuttall FQ. Radioactive method for the assay of glycogen phosphorylases. Anal Biochem 47: 20–27, 1972.

14. Chasiotis D, Sahlin K, and Hultman E. Regulation of glycogenolysis in human muscle at rest and during exercise. J Appl Physiol Respir Environ Exerc Physiol 53: 708–715, 1982.

15. Martin WH, Hoover DJ, Armento SJ, Stock IA, McPherson RK, Danley DE, Stevenson RW, Barrett EJ, and Treadway JL. Discovery of a human liver glycogen phosphorylase inhibitor that lowers blood glucose in vivo. Proc Natl Acad Sci U S A 95: 1776–1781, 1998.

16. Agius L. Role of glycogen phosphorylase in liver glycogen metabolism. Mol Aspects Med 46: 34–45, 2015.

17. Rocha S, Lucas M, Araújo AN, Corvo ML, Fernandes E, and Freitas M. Optimization and Validation of an In Vitro Standardized Glycogen Phosphorylase Activity Assay. Molecules 26: 2021.

18. Hayes JM, Kantsadi AL, and Leonidas DD. Natural products and their derivatives as inhibitors of glycogen phosphorylase: potential treatment for type 2 diabetes. Phytochem Rev 13: 471–498, 2014.

19. Rush JWE, and Spriet LL. Skeletal muscle glycogen phosphorylase akinetics: effects of adenine nucleotides and caffeine. J Appl Physiol 91: 2071–2078, 2001.

20. Cheng K, Zhang P, Liu J, Xie J, and Sun H. Practical Synthesis of Bredemolic Acid, a Natural Inhibitor of Glycogen Phosphorylase. J Nat Prod 71: 1877–1880, 2008.

21. Freeman S, Bartlett JB, Convey G, Hardern I, Teague JL, Loxham SJG, Allen JM, Poucher SM, and Charles AD. Sensitivity of glycogen phosphorylase isoforms to indole site inhibitors is markedly dependent on the activation state of the enzyme. Br J Pharmacol 149: 775–785, 2006.

22. Henke BR, and Sparks SM. Glycogen phosphorylase inhibitors. Mini Rev Med Chem 6: 845–857 2006.

23. Macarthur DG, Seto JT, Raftery JM, Quinlan KG, Huttley GA, Hook JW, Lemckert FA, Kee AJ, Edwards MR, Berman Y, Hardeman EC, Gunning PW, Easteal S, Yang N, and North KN. Loss of ACTN3 gene function alters mouse muscle metabolism and shows evidence of positive selection in humans. Nat Genet 39: 1261–1265, 2007.

24. Haug M, Reischl B, Nübler S, Kiriaev L, Mázala Dag, Houweling PJ, North KN, Friedrich O, and Head SI. Absence of the Z-disc protein α-actinin-3 impairs the mechanical stability of Actn3KO mouse fast-twitch muscle fibres without altering their contractile properties or twitch kinetics. Skeletal Muscle 12: 2022.

25. Kiriaev L, Houweling PJ, North KN, and Head SI. Loss of α-actinin-3 confers protection from eccentric contraction damage in fast-twitch EDL muscles from aged mdx dystrophic mice by reducing pathological fibre branching. Hum Mol Genet 2021.

26. Seto JT, Roeszler KN, Meehan LR, Wood HD, Tiong C, Bek L, Lee SF, Shah M, Quinlan KGR, Gregorevic P, Houweling PJ, and North KN. ACTN3 genotype influences skeletal muscle mass regulation and response to dexamethasone. Science Advances 7: eabg0088, 2021.

27. Wyckelsma VL, Venckunas T, Houweling PJ, Schlittler M, Lauschke VM, Tiong CF, Wood HD, Ivarsson N, Paulauskas H, Eimantas N, Andersson DC, North KN, Brazaitis M, and Westerblad. Loss of α-actinin-3 during human evolution provides superior cold resilience and muscle heat generation. Am J Hum Genet 108: 446–457, 2021.

28. Roberts MD, Young KC, Fox CD, Vann CG, Roberson PA, Osburn SC, Moore JH, Mumford PW, Romero MA, Beck DT, Haun CT, Badisa VLD, Mwashote BM, Ibeanusi V, and Kavazis AN. An optimized procedure for isolation of rodent and human skeletal muscle sarcoplasmic and myofibrillar proteins. J Biol Methods 7: e127, 2020.

29. Blanco A, and Blanco G. Chapter 19 - Integration and Regulation of Metabolism. In: Med Biochem, edited by Blanco A, and Blanco G Academic Press, 2017, p. 425–445.

30. Kasvinsky PJ, Shechosky S, and Fletterick RJ. Synergistic regulation of phosphorylase a by glucose and caffeine. J Biol Chem 253: 9102–9106, 1978.

31. Macarthur DG, Seto JT, Chan S, Quinlan KGR, Raftery JM, Turner N, Nicholson MD, Kee AJ, Hardeman EC, Gunning PW, Cooney GJ, Head SI, Yang N, and North KN. An Actn3 knockout mouse provides mechanistic insights into the association between -actinin-3 deficiency and human athletic performance. Hum Mol Genet 17: 1076–1086, 2008.

32. Zhu Z, Guo Y, Shi H, Liu C-L, Panganiban RA, Chung W, O’Connor LJ, Himes BE, Gazal S, Hasegawa K, Camargo CA, Qi L, Moffatt MF, Hu FB, Lu Q, Cookson WOC, and Liang L. Shared genetic and experimental links between obesity-related traits and asthma subtypes in UK Biobank. Journal of Allergy and Clinical Immunology 145: 537–549, 2020.

